# Fgf23 expression increases atherosclerotic plaque burden in male *ApoE* deficient mice

**DOI:** 10.1101/2024.07.01.601461

**Authors:** Karolina Lindberg, Olga Ovchinnikova, Matthias B. Moor, John Pirault, Daniel FJ Ketelhuth, Hannes Olauson, Göran K. Hansson, Tobias E. Larsson

## Abstract

**Introduction:** Components of both the innate and adaptive immune system impact on arterial walls in atherosclerosis. Fibroblast growth factor-23 (FGF23) is a phosphate regulating hormone linked to cardiovascular disease (CVD) in patients with and without chronic renal disease. However, it remains controversial whether FGF23 is merely a biomarker or contributes to CVD. Here, we overexpressed FGF23 in *ApoE-/-* mice to delineate the role of FGF23 in atherogenesis.

**Methods and Results:** 10-week old *ApoE-/-* mice received a hydrodynamic tail vein with a plasmid encoding for Fgf23, and were sacrificed 10 weeks later. Fgf23 concentrations increased more than 400-fold in the Fgf23 treated group, remaining high throughout the experiment. Mice in the Fgf23 group developed hypophosphatemia, secondary hyperparathyroidism and a moderate increase in plasma creatinine concentrations. Male *ApoE-/-* mice exposed to high Fgf23 developed larger atherosclerotic lesions compared to controls, in two different locations of aorta, whereas no differences in plaque burden were seen between female *ApoE-/-* mice and controls. Serum IL-6 concentrations were increased in the Fgf23 group, associated with a vascular inflammatory response of recruited macrophages and neutrophils, and with a shift of CD4+ T effector cells from Th1 to Th17 and migration of lymphocytes to the spleen.

**Conclusion:** Fgf23 increases the atherosclerotic burden in male *ApoE-/-* mice and alters both the innate immune system and T cell subpopulations, generating an inflammatory environment that may promote adverse clinical outcomes associated with Fgf23 excess.

## Introduction

Atherosclerosis is a disease of the arterial wall with multiple etiologies, including an inflammatory component mediated both by the innate and adaptive immune system. Retention of lipoproteins, particular low-density lipoproteins (LDL) and cholesterol in the intimal layer of the arterial wall promotes endothelial cells to express adhesion molecules and chemokines on their surface, leading to recruitment of immune cells to the subendothelial space. Infiltration of the vessel wall with monocyte/macrophages and T lymphocytes is a hallmark of atherosclerosis and contributes to growth of atherosclerotic plaques. The proinflammatory environment in the plaque and proteolytic enzymes create a necrotic core, weakens the surrounding collagenous matrix and, eventually, disrupts the plaque. Plaque rupture and subsequent thromboembolic occlusion are, together with plaque erosion, the most frequent mechanisms causing lethal complications of atherosclerosis such as myocardial infarction and ischemic stroke (1,2).

Fibroblast growth factor-23 (FGF23) is a bone-derived hormone that promotes phosphaturia and 1,25(OH)2-vitamin D metabolism. Circulating FGF23 concentrations are inversely correlated to renal function and are recognized as a potential uremic toxin that promotes cardiovascular disease. FGF23 concentrations correlate with endothelial dysfunction, activation of the renin angiotensin aldosterone system, heart failure and adverse clinical outcomes across a broad range of renal function including patients on dialysis treatment (3–8).

Several epidemiological studies revealed that elevated circulating FGF23 concentrations are associated with endothelial dysfunction, left ventricular hypertrophy (LVH) and mortality. (3,9–11). Controversies however remain if FGF23 is predominantly a marker or rather a cause of disease, or under which circumstances the benefits in reducing systemic phosphate load may outweigh the cardiovascular risk of elevated Fgf23 concentrations (12,13). Several mechanisms linking excess FGF23 to cardiovascular disease were postulated including indirect actions (e.g. abnormal mineral metabolism such as vitamin D deficiency and hyperparathyroidism) and direct actions (e.g. cardiac hypertrophic effects via FGF-receptor signaling) (14,15).

Herein we systemically overexpressed FGF23 in mice prone to spontaneous development of atherosclerosis (apolipoprotein E-deficient, *ApoE^-/-^* mice) to delineate the role of FGF23 in atherogenesis.

## Material and Methods

### Fgf23-containing plasmid

A pcDNA3.1/ V5-His-TOPO plasmid (Invitrogen) containing the full-length rat *Fgf23* sequence with a stabilizing mutation (R176Q) under control of a CMV promoter was kindly provided by Dr Susan Schiavi, at the time at Genzyme, MA, USA.

### Animal experiments

*ApoE^-/-^* mice on C57BL/6J background from in-house breeding at Karolinska University Hospital in Stockholm were used in this study. Ten weeks old mice received a single hydrodynamic tail vein injection (16) with either 10 μg Fgf23- containing plasmid in saline (Fgf23 group) or saline only (control group). The total volume (1/10 of the body weight) was rapidly injected in <10 s. Mice were sacrificed at 20 weeks of age. In total, 62 mice were injected and eligible for final analysis: 34 males (n=15 in Fgf23 group and n=19 in control group) and 28 females (n=13 in Fgf23 group and n=15 in control group). The local ethics committee approved all animal experiments.

### Serum biochemistry

Blood was collected before the hydrodynamic tail vein injection, and at weeks 1, 3, 5, 8 and 10 after injection. At sacrifice, blood cells were counted in the whole mouse blood on ABC Vet Animal Blood Counter (ABX Hematologie, Montpellier, France). Serum concentrations of calcium, phosphate, and creatinine were measured using quantitative colorimetric assay kits (BioAssay; BioChain Institute, Inc., Newark, CA). Serum parathyroid hormone (PTH) was measured using a mouse 1-84 PTH ELISA (Immutopics, San Clemente, CA), FGF23 was measured using the mouse FGF23 (c-term) ELISA (Immutopics) and IL-6 – using mouse interleukin-6 (IL-6) ELISA (BioLegend, CA, US). Serum 1,25-Dihydroxyvitamin D (1,25(OH)2D) was measured using an EIA assay (Immunodiagnostic Systems, Scottsdale, AZ). Cholesterol and triglycerides were measured by enzymatic colorimetric kits (Randox Lab. Ltd. Crumin, UK).

### Histological analysis of atherosclerotic lesions

Hearts and ascending aorta were dissected immediately upon sacrifice and preserved for immunohistochemistry and analysis of lesions. Cryosections and analysis of lesions were performed as described (17). Briefly, hearts were serially sectioned from the proximal 1 mm of the aortic root on a cryostat. Hematoxylin- and oil red O-stained sections were used to evaluate lesion size on 8 hematoxylin- and Oil Red O-stained sections, collected at every 100 μm over a 1 mm segment of the aortic root. For each section, images were captured for measurement of the surface areas of the lesion(s) and of the lumen of entire vessel.

*En face* lipid accumulation was determined in the aortic arch from mice by using Sudan IV staining. Briefly, dissected arches were fixed in 4% neutral buffered formalin. Samples were then cut longitudinally, splayed, pinned and subjected to Sudan IV staining. The area covered by plaques in each aortic arch was calculated as a percent of the total surface area of the arch, excluding branching vessels.

Picrosirius red staining was used for the assessment of collagen fibers in the lesions. All sections were analyzed under linear polarized light at ×20 magnification on a Leica DMRB microscope. The total collagen content of lesions was measured using Leica QWin and calculated as the ratio of thresholded chromogen area to lesion area.

### Gene expression

Total RNA was extracted from aortas and analyzed as previously described(18). Premanufactured primers and probe sets (Assay-on-demand, Applied Biosystems) for *Cd68*, *Cd3*, *Mpo*, *Mcp1*, *Cxcl1*, *Ccr6*, *Cxcr6*, *T-bet*, *Gata3*, *Foxp3*, *Ror*γ*t*, *Ifn-*γ, *Tgf-*β, *Il10*, *Tnf-*α, *Col1a1*, *Lox*, *Mmp9*, *Mmp13*, *RunX2* and a house keeping gene (hypoxanthine-guanine phosphoribosyltransferase, *Hprt*) were used to perform quantitative real time PCR (rtPCR). Data were analyzed with the 2^−ΔΔ^ CT method (Applied Biosystems). Analysis of Klotho mRNA expression in renal and aortic tissues and Fgfr1 mRNA in aortic tissue using SYBRGreen® method (BioRad, CA, US) and β-actin as reference gene.

### Immunostaining of the atherosclerotic lesions

Primary antibodies included rat anti-mouse CD68 (Serotec Ltd., Oxford, United Kingdom), rat anti-mouse VCAM-1, biotinylated mouse anti-mouse I-Ab (both from Pharmingen, San Diego, CA, USA), rat anti-mouse CD4, rat anti-mouse Gr1/Ly6G (both from BD Biosciences, NJ, USA), or rabbit anti–α-smooth muscle actin (Abcam, Cambridge, UK). The binding of primary antibody on consecutive sections was detected using ABC kit/Peroxidase and visualized with Vector NovoRed or Vector DAB substrate kit (Vector laboratories).

### Gene expression of peritoneal macrophages

Male *ApoE^-/-^* mice injected with Fgf23 plasmid or saline were used for peritoneal lavage. Briefly, 5 mL of PBS was injected in the abdomen and harvested after peritoneal lavage. Cells were washed and seeded on 6 well plates for 3h at 37 C. Floating cells were removed followed by 3 washes with PBS. Adherent cells were kept in complete RPMI medium. After 48h cells were harvested for mRNA extraction and gene expression analysis.

### Analysis of serum antibodies to oxidized LDL

For analysis of circulating antibodies to oxidized LDL (oxLDL), 50 µL of oxidized LDL (oxLDL) (10 µg/ml in PBS pH 7.4) was added to 96-well ELISA plates and incubated overnight at 4°C. Coated plates were washed with PBS and blocked with 1% gelatin (Gibco Invitrogen, Carlsbad, CA, USA) in PBS for 1h at room temperature. Next, plates were washed and incubated for 2h with mouse plasma and diluted in Tris-buffered saline (TBS)/gelatin 0.1%. After washing, total IgG levels were measured using enzyme-conjugated anti-mouse antibodies (BD Biosciences, Franklin Lakes, NJ, USA). The plates were washed, and colorimetric reactions were developed with TMB (BD Biosciences, Franklin Lakes, NJ, USA). The absorbance was measured on a microplate reader (VersaMax, Molecular Devices, Sunnyvale, CA, USA).

### Statistical analysis

All values are expressed as mean+SEM unless otherwise indicated. Nonparametric Mann–Whitney U test was used for comparisons between two groups. Differences were considered significant at P-values < 0.05. All statistical analyses were performed using GraphPad Prism for Mac OS X (GraphPad Software, Inc., CA, USA).

## Results

### Fgf23 concentrations and mineral metabolism

First, we assessed the efficacy and physiological consequences of Fgf23 gene delivery. Baseline plasma Fgf23 concentrations were similar between the treatment groups (control vs Fgf23; 175 ± 28.16 vs 159.3 ± 21.46 pg/mL, *P*=0.84). After hydrodynamic tail vein injection, Fgf23 levels remained low in control mice at all time points whereas in Fgf23 mice, Fgf23 concentrations rose after 1 week (>400 fold increase), peaking at over 57 000 pg/ml on week 3 after the injection, and remaining high until sacrifice. (Figure 1A).

**Figure 1.**
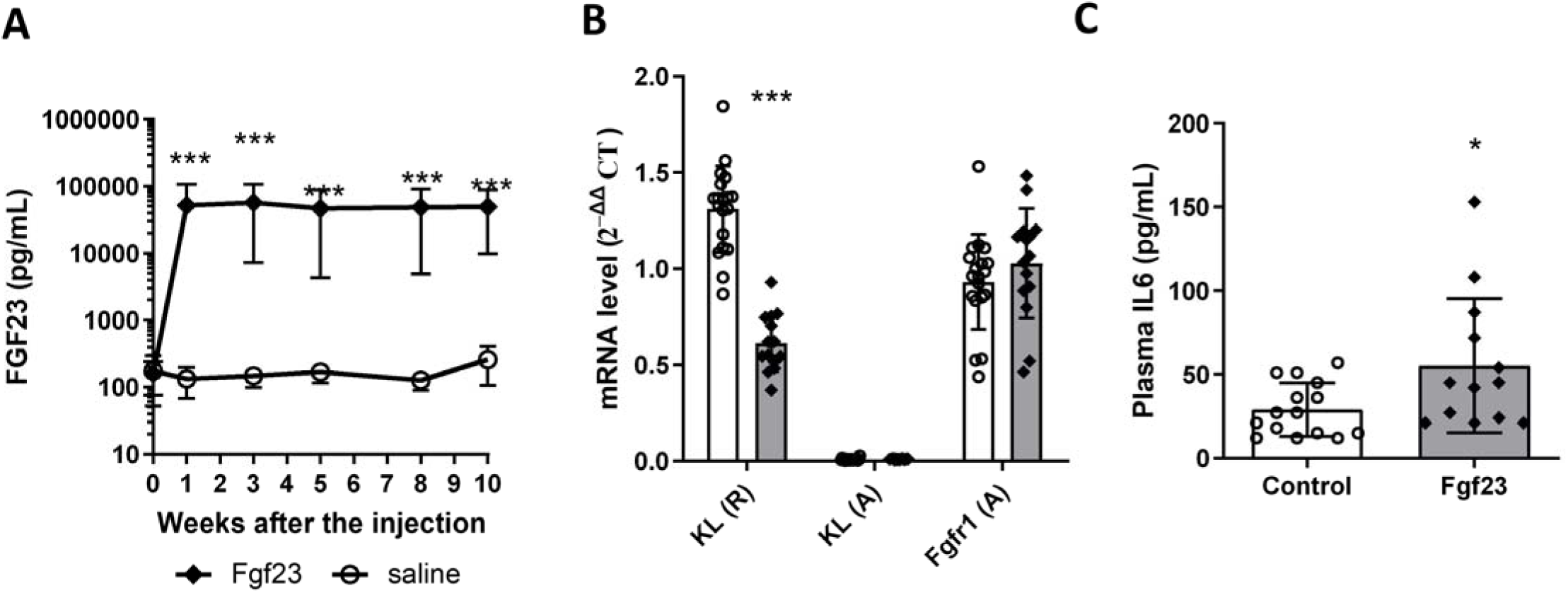
(A) Plasma FGF23 concentrations were analyzed at baseline (week 0), at weeks 1, 3, 5, and 8 after the injection, and at the endpoint (10 weeks after the injection) to monitor efficiency of Fgf23 expression in *ApoE-/-* male mice. (B) Analysis of Klotho mRNA in aortic and renal extracts and Fgfr1 mRNA in aortic extracts of *ApoE-/-* male mice. Data are presented as 2-ΔΔCt value of a specific gene normalized to RNA of a house-keeping gene (β-actin). (C) Plasma concentration of IL6 at the endpoint of the experiment. * *P*<0.05; *** *P*<0.001; n of mice = 19 and 15 for control and Fgf23 groups, respectively. Error bars indicate standard deviation.

Serum phosphate concentration was lower in the Fgf23 group than in controls, consistent with the phosphaturic effect of Fgf23. Fgf23 mice also developed evidence of secondary hyperparathyroidism with lower calcium levels and higher parathyroid hormone (PTH). Renal function inferred from serum creatinine concentrations was similar between groups initially but at sacrifice the Fgf23 group had elevated creatinine concentrations. No difference in serum urea concentration was observed (Table 1). Body weight at baseline was similar between the treatment groups (control vs Fgf23; 26.14±0.56 and 25.97±0.73 g, *P*=0.85), but at the end of study weight was lower in the Fgf23 group (Table 1)

**Table 1.**
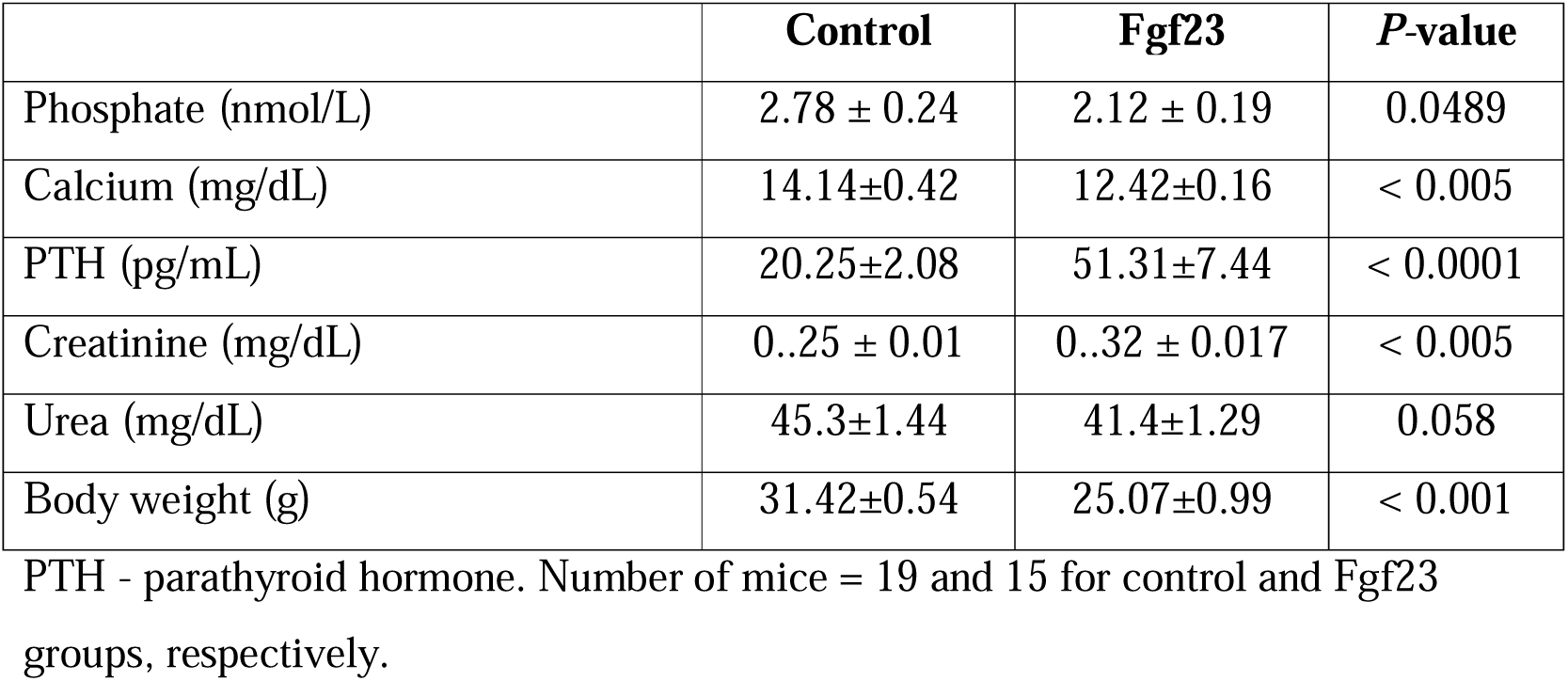
Serum parameters and body weight at the endpoint.

|  | **Control** | **Fgf23** | ***P*-value** |
| --- | --- | --- | --- |
| Phosphate (nmol/L) | 2.78 ± 0.24 | 2.12 ± 0.19 | 0.0489 |
| Calcium (mg/dL) | 14.14±0.42 | 12.42±0.16 | < 0.005 |
| PTH (pg/mL) | 20.25±2.08 | 51.31±7.44 | < 0.0001 |
| Creatinine (mg/dL) | 0.25 ± 0.01 | 0.32 ± 0.017 | < 0.005 |
| Urea (mg/dL) | 45.3±1.44 | 41.4±1.29 | 0.058 |
| Body weight (g) | 31.42±0.54 | 25.07±0.99 | < 0.001 |

PTH - parathyroid hormone. Number of mice = 19 and 15 for control and Fgf23 groups, respectively.

### Fgf23 overexpression promotes atherosclerotic burden in ApoE^-/-^mice

Next, we assessed the effect of *Fgf23* delivery on the vasculature. Male *ApoE^-/-^* mice of the Fgf23 group developed larger atherosclerotic lesions than controls in both the aortic arch and in the aortic root (Figure 2). The diameter of the aortic roots remained similar between groups (Supplementary Figure 1).

**Figure 2.**
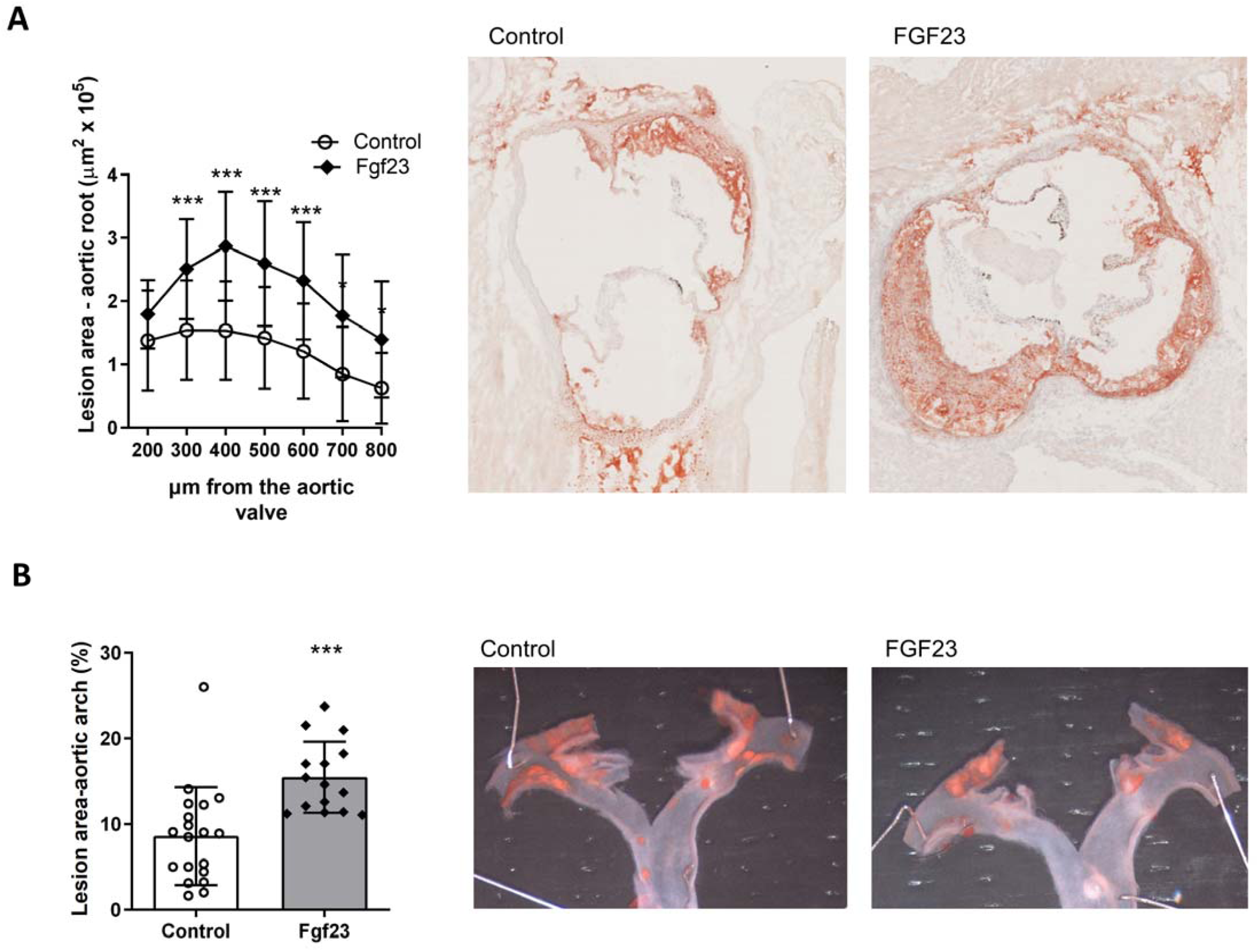
*ApoE-/-* male mice received a single hydrodynamics-based transfection of Fgf23-containing plasmid. Overexpression of FGF23 increased atherosclerotic lesion size male mice. (A) Aortic roots were stained with Oil Red O. The absolute lesion area was calculated at 7 levels above the aortic root. Representative micrographs of mouse aortic roots; magnification x50. (B) En face lesion area in the aortic arch. Graphs represent the percent (%) of Sudan IV stained area. * *P*<0.05, *** *P*<0.001; n of mice = 19 and 15 for control and Fgf23 groups, respectively. Error bars indicate standard deviation.

The atherosclerotic lesions appeared larger in female mice than in males, however no significant differences in lesion size were observed between the treatment groups in females (Supplementary Figure 2). All further analyses were therefore performed in males only.

As one of the hallmarks of atherosclerosis is a pro-inflammatory milieu, we assessed serum IL-6 concentrations. IL-6 was increased in the Fgf23 group compared to controls (Figure 1B).

Plasma total cholesterol did not differ significantly between the two experimental groups, whereas plasma triglycerides level was lower in the plasmid injected mice (Supplementary Figure 3). Titers of atheroprotective antibodies against oxLDL were comparable between experimental groups (data not shown). To summarize, male *ApoE-/-* mice exposed to high Fgf23 had a significantly higher atherosclerotic plaque burden in two aorta sites and an increased proinflammatory milieu.

### Gene expression

Aortic *Fgfr1* mRNA was detected at similar levels in both treatment groups, whereas mRNA levels of renal FGF23 co-receptor *Klotho* were almost undetectable. This suggests that the effect of FGF23 on atherosclerotic plaque burden in the Fgf23 group is independent of vascular Fgf23-Klotho signaling. Consistent with our previous results (19), *Klotho* mRNA expression in kidney was suppressed in Fgf23 mice (Figure 1B).

Next, we aimed to gain insight in the regulation and consequences of aortic inflammation. Aortic transcript levels of macrophage marker *Cd68*, neutrophil marker *Mpo* and monocyte attraction factors *Mcp1* and *Cxcl1* were increased in the Fgf23 group. Transcripts related to T-cell homing (*Ccr6* and *Cxcr6*) and osteoblast differentiation (*Runx2*) did not differ between the groups (Table 2).

**Table 2.**
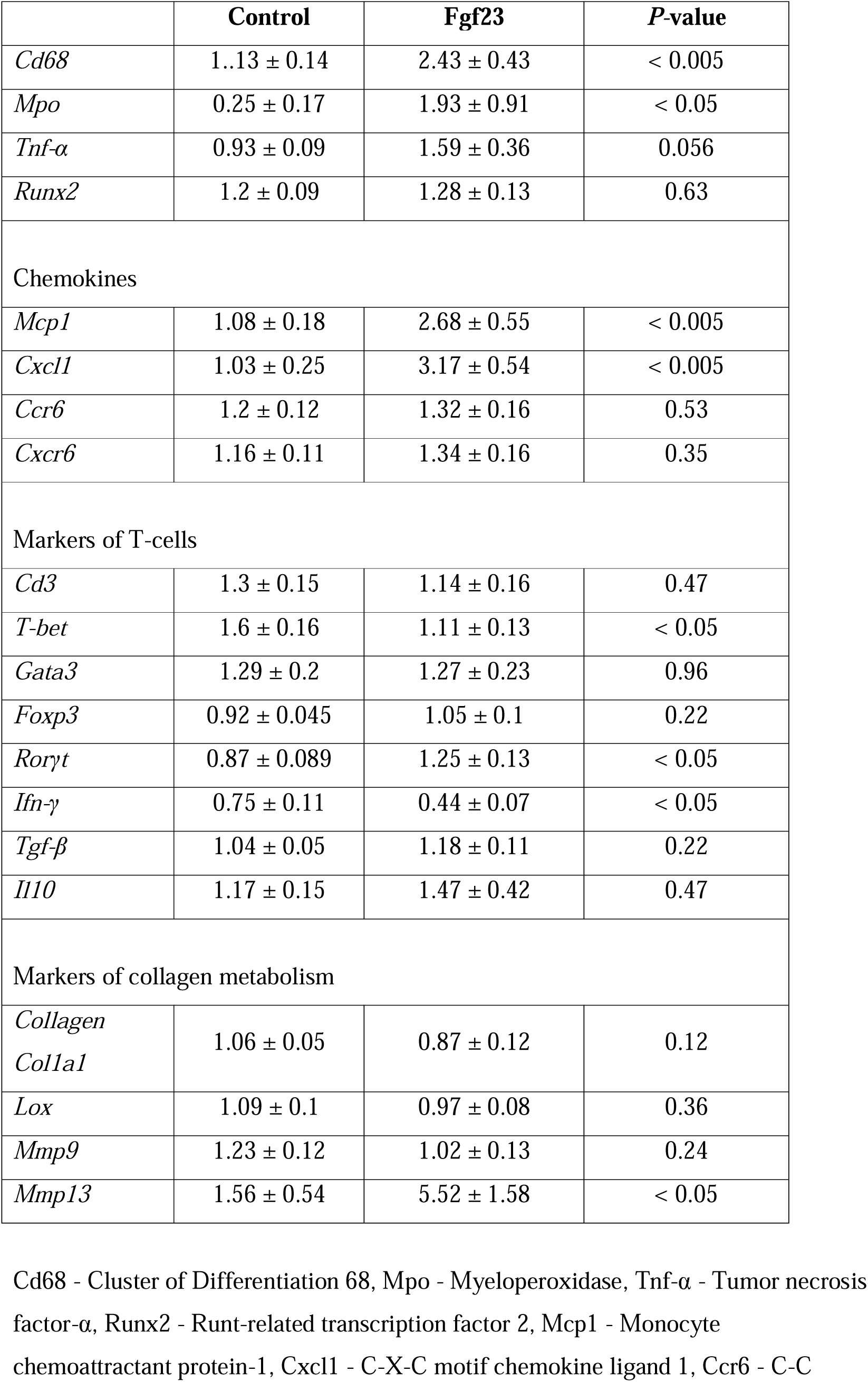
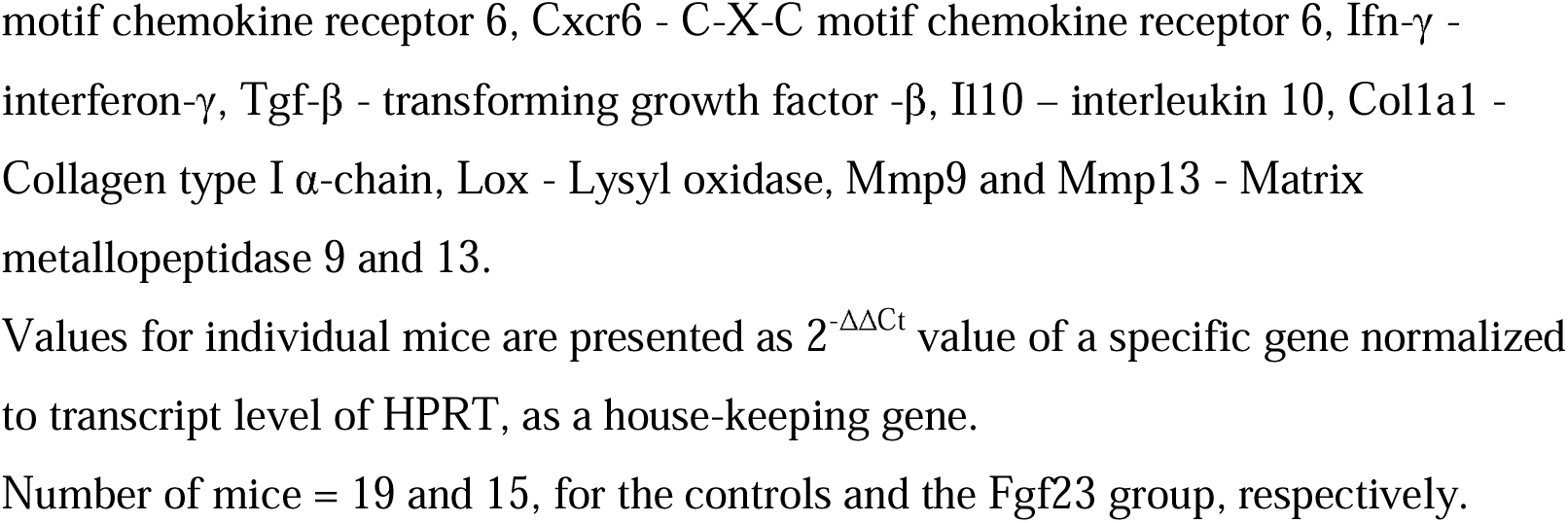
mRNA expression in aortae of *ApoE*-/- male mice.

Transcripts of general T cell marker *Cd3* were similar between groups, but *T-bet* and *Ifn-*γ associated with Th1 cells and a major cytokine of Th1 cells, were decreased upon Fgf23 overexpression. Conversely, Th17 cell signature assessed by *Ror*γ*t* was increased upon FGF23. *Gata3* and *Foxp3* associated with Th2 cells and Treg cells, respectively, and *Tgf-*β and *Il10* were comparable between groups (Table 2). As a consequence of inflammation, matrix metalloproteinase (*Mmp) 13* but not *Mmp-9* was elevated in aortae from Fgf23 mice. Transcripts of *Col1a1,* encoding collagen type 1alpha1 and *Lox* encoding a collagen stabilizing enzyme were all comparable between groups (Table 2). To summarize, mice in the Fgf23 group presented distinct transcriptional changes in the aorta consistent with an increased presence or activity of macrophages, neutrophils, a T cell shift from Th1 towards the Th17 subpopulation, and increased transcripts among chemokines and proteases.

### Phenotype of atherosclerotic plaques in mice overexpressing Fgf23

To substantiate the findings from the gene expression studies we next assessed the corresponding protein level. CD68+ macrophage and Gr1/Ly6G+ neutrophil infiltration in the intima of the vessel wall were higher in the Fgf23 group compared to controls (Figure 3A-B). An increased CD68+ macrophage infiltration was also present in the adventitia (Figure 3A). Protein staining for vascular cell adhesion molecule-1 (VCAM-1), which is pivotal in leukocyte infiltration and lesion progression, was also increased in the Fgf23 group (Figure 3C). However, the CD68 and VCAM-1 signals revealed no qualitative differences in plaque cell composition between groups when they were normalized to the lesion size (Table 3). The lesions in the aortic roots of Fgf23 mice did not differ significantly from those of control mice with regard to necrotic core volume or the number of α-actin+ smooth muscle cells in the fibrous caps (data not shown).

**Figure 3.**
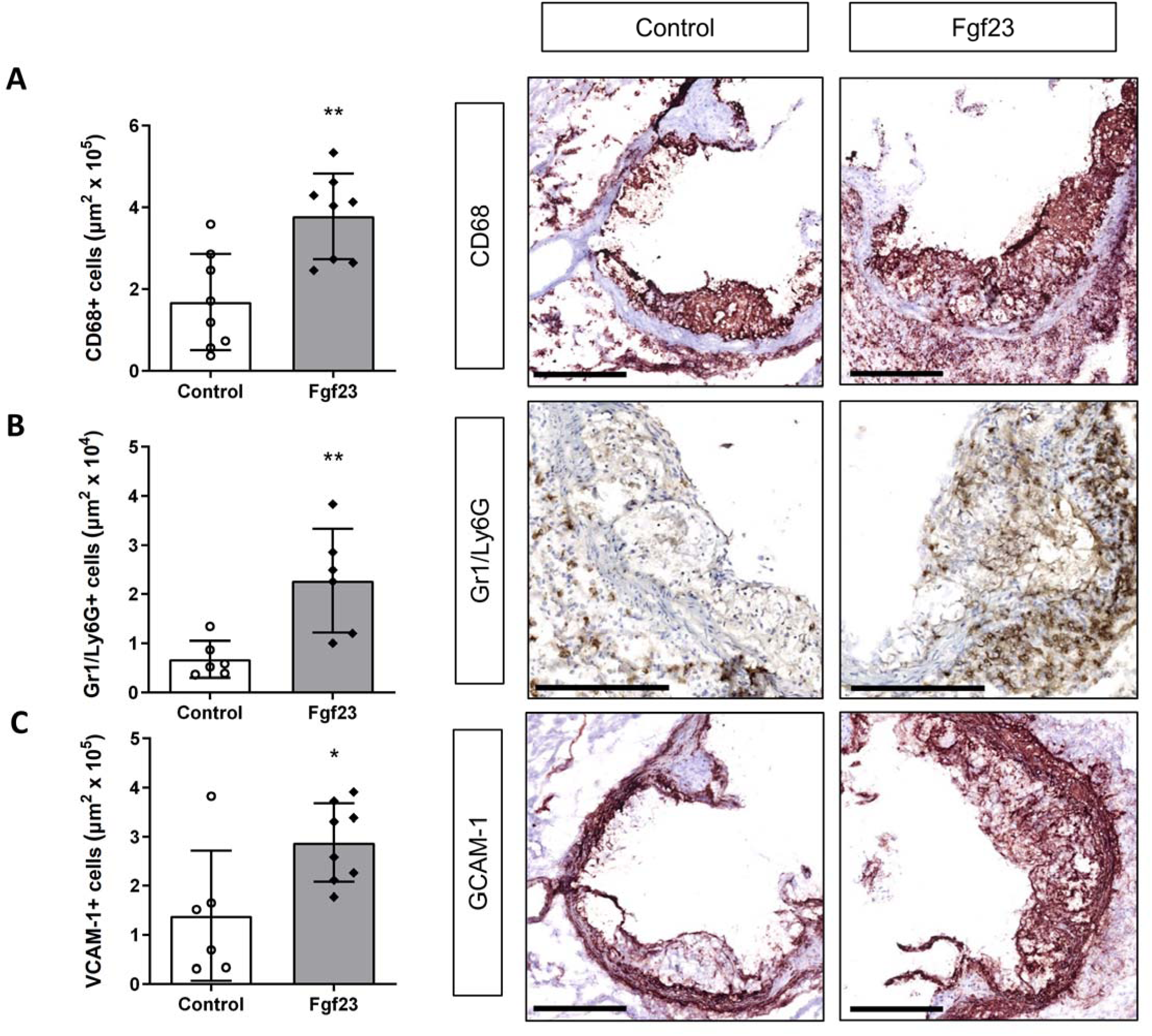
Fgf23 overexpression induced CD68+ macrophage (A) and Gr1/Ly6G+ neutrophils infiltration (B), as well as VCAM-1 expression (C) in vessel wall of *ApoE*−/− male mice. Right panels show representative micrographs from each group. (A) Black arrows point to the more intensive CD68+ staining in adventitia. \**P* < 0.05; \*\**P* < 0.01; n= 6-8 per group. Error bars indicate standard deviation.

**Table 3.** Plaque composition at the endpoint.

|  | Control | Fgf23 | *P*-value |
| --- | --- | --- | --- |
| CD68+ cells (% stained area) | 28.8 ± 4.7 | 34.0 ± 2.4 | 0.34 |
| Gr1/Ly6G+ cells (% stained area) | 0.74 ± 0.14 | 2.9 ± 0.65 | < 0.005 |
| VCAM-1+cells (% stained area) | 22 ± 4.9 | 28.1 ± 2.3 | 0.23 |
| CD4+ cells (% stained area) | 4.1 ± 1.5 | 3.2 ± 1.1 | 0.39 |
| I-Ab+ cells (% stained area) | 3.5 ± 0.83 | 1.65 ± 0.13 | < 0.05 |
| α-actin+ cells (stained area in fibrous cap, μm<sup>2</sup> x 10<sup>3</sup>) | 16.6 ± 5.5 | 24.7 ± 6.3 | 0.5 |
| Necrosis area (% stained area) | 9.5 ± 2.6 | 8.3 ± 2.2 | 0.76 |

Activated Th1 cells induce expression of the major histocompatibility complex (MHC) class II molecules (I-Ab antigen) on the cells in the plaques. Consistent with the transcriptional changes reported above, I-Ab+ staining was diminished in Fgf23 mice compared to controls (Table 3). The number of overall CD4+ T-lymphocytes in intima and adventitia were unchanged in the Fgf23 group (data not shown). Overall, the immunohistochemistry analyses revealed a relative decrease in a Th1 cell marker, but all other differences between the groups depended on atherosclerotic plaque size.

### Fgf23 expression promotes macrophage activation

To further describe the immunological mechanisms of the increased atherosclerotic burden in mice overexpressing Fgf23, separate groups of male *ApoE^-/-^*mice were injected with either an Fgf23 containing plasmid or saline, followed by isolation and analysis of peritoneal macrophages. Transcripts *Mcp1* and *Cxcl1* were increased in peritoneal macrophages from mice in the Fgf23 group (Figure 4). Transcripts of *Tnf-*α and *Chil3*, markers of M1 and M2-macrophages, respectively, were elevated in peritoneal macrophages in mice from the Fgf23 group (Figure 4). However, expression of *Mmp13* in peritoneal macrophages was similar between groups (Figure 4). Collectively, the peritoneal analysis revealed a general increase in macrophage markers but no signs of increased protease expression.

**Figure 4.**
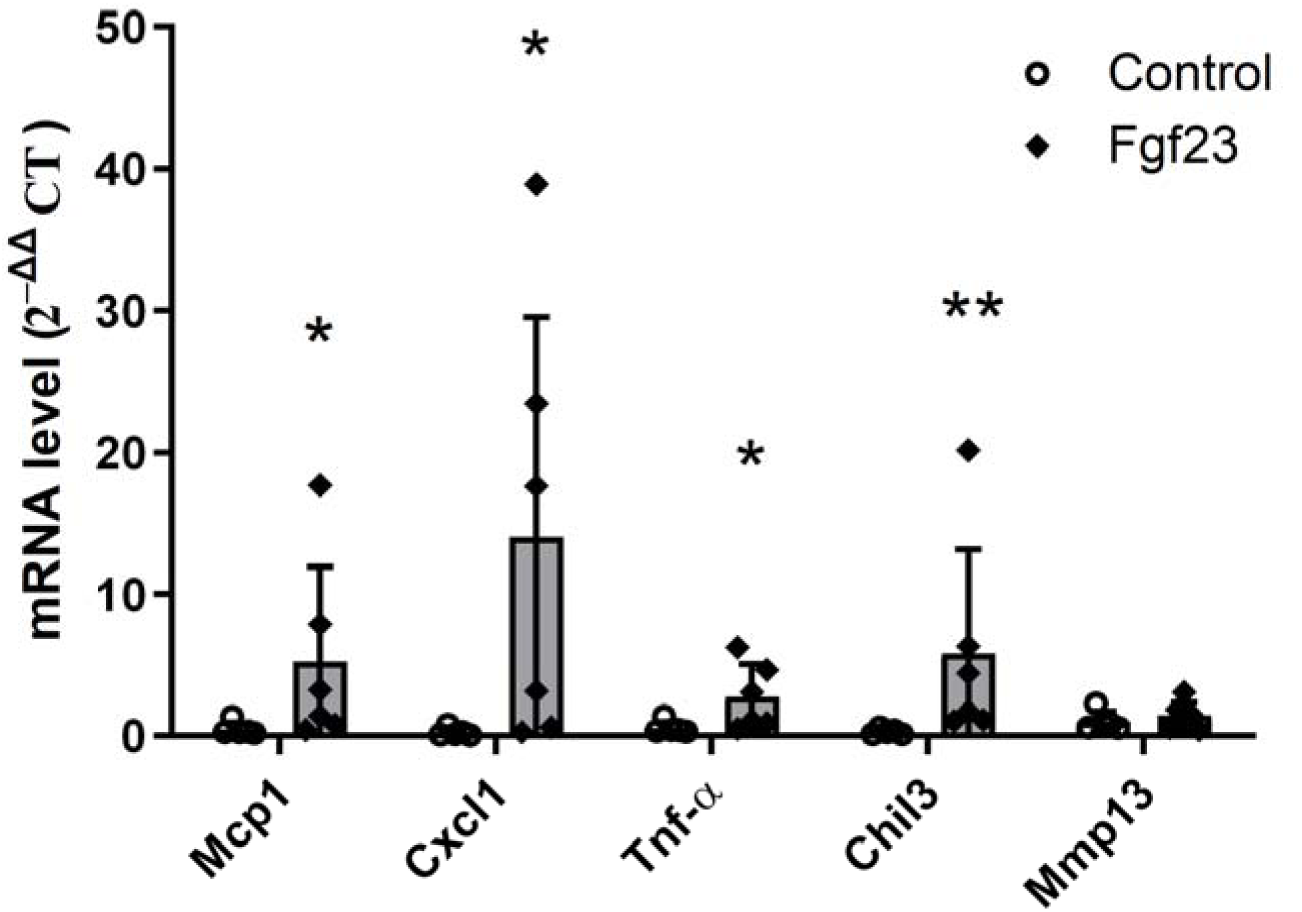
Separate group of *ApoE-/-* male mice were injected with Fgf23-containing plasmid or saline. mRNA expression of Mcp1, Cxcl1 and markers of macrophages activation was analyzed in peritoneal macrophages. Values for individual mice are presented as 2^-ΔΔCt^ value of a specific gene normalized to RNA of a house-keeping gene (HPRT). \**P* < 0.05; \*\**P* < 0.01, n – 5-6 per group. Error bars indicate standard deviation.

### Lymphocyte population in blood and spleen

Circulating immune cells play a major role in initiation and progression of atherosclerosis. Therefore, the effects of Fgf23 overexpression on circulating blood cells were analyzed in whole blood. Mice overexpressing Fgf23 showed significantly reduced absolute numbers of circulating lymphocytes, whereas monocytes and granulocytes were comparable (Supplementary Figure 4A). Circulating erythrocytes and platelets were also comparable between groups (data not shown).

To examine the distribution of cell populations in the spleen and paraaortic lymph nodes, FACS analysis was performed on mouse spleens and lymph nodes harvested at time of sacrifice. The number of CD4+ and CD8+ cells in spleen was significantly increased in the Fgf23 group. No significant differences were found in monocyte/macrophage populations in the spleen or in any of the cell populations of paraaortic lymph nodes (Supplementary Figure 4). To summarize these findings, we found that Fgf23 overexpression promoted a decrease in circulating but an increase in splenic lymphocytes.

## Discussion

In the present study we set out to delineate the role of Fgf23 in atherosclerotic plaque formation and in the immune cell landscape involved in atherogenesis. We demonstrate that chronic exposure to Fgf23 exacerbates atherosclerotic plaque burden in male *ApoE^-/-^* mice. This finding was associated with a local inflammatory response including enhanced recruitment of macrophages and neutrophils into the subendothelial space and a T cell shift from Th1 to the Th17 subpopulation but also with a tendency to splenic lymphocyte homing, accompanied by lymphocytopenia in peripheral blood.

A plethora of data in both human patients and experimental models have demonstrated that excessive circulating Fgf23 and phosphate concentrations are associated with impaired cardiovascular health (3,20,21). The potential causes include vascular calcification, endothelial damage, Klotho and vitamin D deficiency, and RAAS activation by Fgf23 and sodium retention (4,22). Direct vascular effects of Fgf23 are largely unclear, as Fgf23 signaling is unlikely to occur in the absence of Klotho expression. However, via Klotho-independent signaling in the liver, Fgf23 excess can promote an inflammatory milieu by inducing expression of the pro-inflammatory cytokine IL-6 (23,24), which could adversely affect the vasculature. In addition, the present study showed that Klotho downregulation in the kidney is a lasting consequence of chronic Fgf23 excess. Klotho deficiency alone has previously been reported as a biomarker for atherosclerosis (25,26), and it may have direct or indirect effects on atherogenesis.

The pro-atherosclerotic effect of FGF23 is consistent with clinical data demonstrating an association between circulating FGF23 and increased carotid intima media thickness (IMT) and plaque size in CKD patients (27). In the PIVUS study, FGF23 was associated with luminal stenosis, a surrogate for atherosclerosis, in most vascular territories, as measured by whole-body MRI angiography (9).

Adherence of classical monocytes to the endothelium and transmigration into the subendothelial space is a well-defined first essential step in atherosclerotic lesion development (28). Our study demonstrated that overexpression of Fgf23 *in vivo* modulated the aortic expression of vascular adhesion molecule, leading to increased expression of the macrophage specific chemokines MCP1 and CXCL1 and macrophage infiltration in the vessel wall.

The present data also confirm that circulating IL-6 concentration was elevated in mice with chronic Fgf23 excess, in line with previous studies of the hepatic response to Fgf23 (24). IL-6 is released from activated monocytes and macrophages as well as vascular cells in response to proinflammatory cytokines such as IL-1 (2,29). *In vitro,* FGF23 increases the number of macrophages and augments their inflammatory response in culture (30). Similarly, FGF23 was positively associated with markers of endothelial cell injury, including VCAM-1, in kidney transplant recipients (31).

CD68 and VCAM-1 expression were increased in the Fgf23 group, proportional to the size of atherosclerotic lesions. In these animals, elevated CD68+ staining was found not only in intima but also in tissue adjacent to the aorta, suggesting increased degree of adventitial inflammation and consequently protease activity (32–34).

Notably, Fgf23 transgenic mice are characterized by increased bone matrix degradation due to increased protease secretion (35), which could also be paralleled by vascular matrix remodeling. In agreement, the aortic transcripts of the major collagenase detected in mice, *Mmp13*, were increased in the Fgf23 group. This was paralleled by lower amount of collagen fibers in the aortic plaque lesions, suggesting that high systemic FGF23 plays a role in vascular matrix remodeling and the transition from early to advanced inflamed atherosclerotic lesions.

Finally, Fgf23 overexpression diminished the abundance of Th1-associated transcriptional and protein markers, in favor of increased Th17-associated gene expression. The reduction of Th1 markers was paralleled by reduced expression of the MHC-II protein, I-A^b^, which is induced by interferon-γ. This may imply a reduction in the capacity for antigen presentation. The concomitant increase in Th17 markers was accompanied by neutrophil markers. Interestingly, IL-17A, the signature cytokine of Th17 cells, is known to enhance neutrophil activation and the formation of neutrophil extracellular traps that may contribute to atherosclerosis. (36). It remains to be determined whether such modulation of immune activity and inflammation could impact on host defense against pathogens leading to recurrent infections, a common cause of death in patients with CKD (37).

This study differs from previously published studies in some key respects. For instance, excessive circulating Fgf23 concentrations in the *Hyp* mouse did not directly lead to cardiac disease in the absence of hyperphosphatemia (38). Importantly, in the present study atherosclerotic burden was increased by Fgf23 even in a state of hypophosphatemia, and without any changes in plasma LDL concentrations. The reason may be a different experimental readout, and an experimental model already prone to atherosclerosis. Next, the chronic Fgf23 exposure in our study yielded different results than the report of an acute Fgf23 treatment that inhibited neutrophil activation, adhesion and transendothelial migration in response to acute bacterial infection in a CKD model (39). However, in contrast to a 24 h exposure with Fgf23, our 10-weeks Fgf23 treatment allows time for numerous changes including immune cell recruiting and their potential consequences (i.e. atherogenesis).

This study contains some limitations. First, we used saline in controls instead of injecting a plasmid encoding a non-expressed transcript. However, the observed phenotypic changes in Fgf23 mice correspond well with the expected Fgf23 bioactivity as the main physiological difference between groups. Also, in a previous study of the atherogenic process in *ApoE^-/-^* mice, there were no differences in plaque formation between mice receiving a control vector and PBS (40). Next, this study was not sufficiently powered for a sex-stratified analysis of all results and after the initial experiment, all subsequent experiments were restricted to males. Our findings may point towards a stronger importance of inflammation in the pathogenesis of atherosclerotic plaques in males than females, as suggested before (41). However, it would be important to perform sex-stratified analyses in the future, as sex influences the pathophysiology of inflammation in atherosclerosis, and female *ApoE^-/-^* mice have a larger plaque burden than males at baseline (42,43).

Finally, this study captures the combination of direct Fgf23 effects and indirect effects of disturbances in calcium, phosphate, Klotho, PTH and a potential decrease in renal function, without being able to discriminate the direct from the indirect Fgf23 effects.

In conclusion, we demonstrate that chronic FGF23 excess creates a proinflammatory milieu involving both innate and adaptive immune responses, and that chronic FGF23 excess exacerbates atherosclerosis in a well-studied model organism. These experimental data unravel an additional mechanism to explain how FGF23 may contribute to excess cardiovascular morbidity and mortality in humans, calling for an evaluation of immunomodulation to improve cardiovascular outcomes in CKD.

## Acknowledgements

We thank Ingrid Törnberg and André Strodthoff for technical assistance. We thank the AKM Animal Facility staff for help with animal care, and especially Kristina Edvardsson for the help with hydrodynamic tail vein injections.

## Conflicts of interest

TL is employed by Guard Therapeutics, Stockholm, Sweden. All authors have no other conflicts to declare.

## Funding

This study was supported by the Swedish Research Council (grant no 01789), Swedish Kidney Foundation, Swedish Heart-Lung Foundation, grants from Karolinska Institutet and Karolinska University Hospital to TL; the Stockholm County Council – ALF to GKH; the Foundation for Geriatric Diseases at Karolinska Institutet to OO; CIMED to HO. MBM was supported by the Swiss National Science Foundation (grant number 214187).

## Supplemental data

**Supplemental Figure 1.**
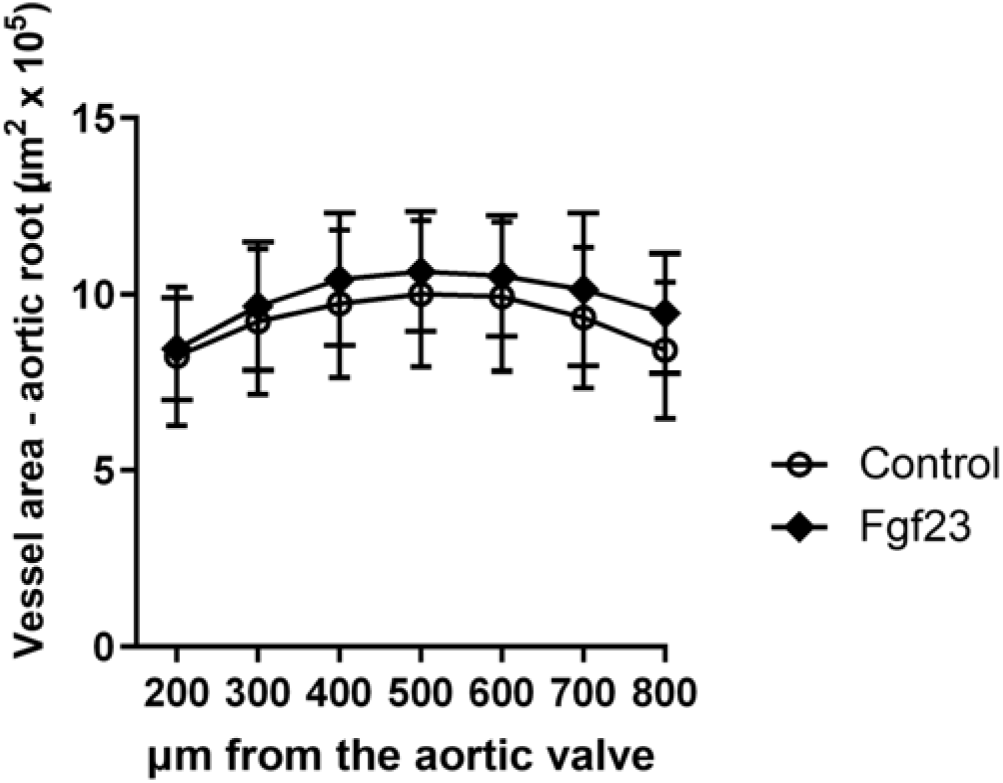
Overexpression of FGF23 did not influence vessel cross-sectional area in aortic root in *Apo£-/-* male mice. The absolute vessel area was calculated at 7 levels above the aortic root. Error bars indicate standard deviation.

**Supplemental Figure 2.**
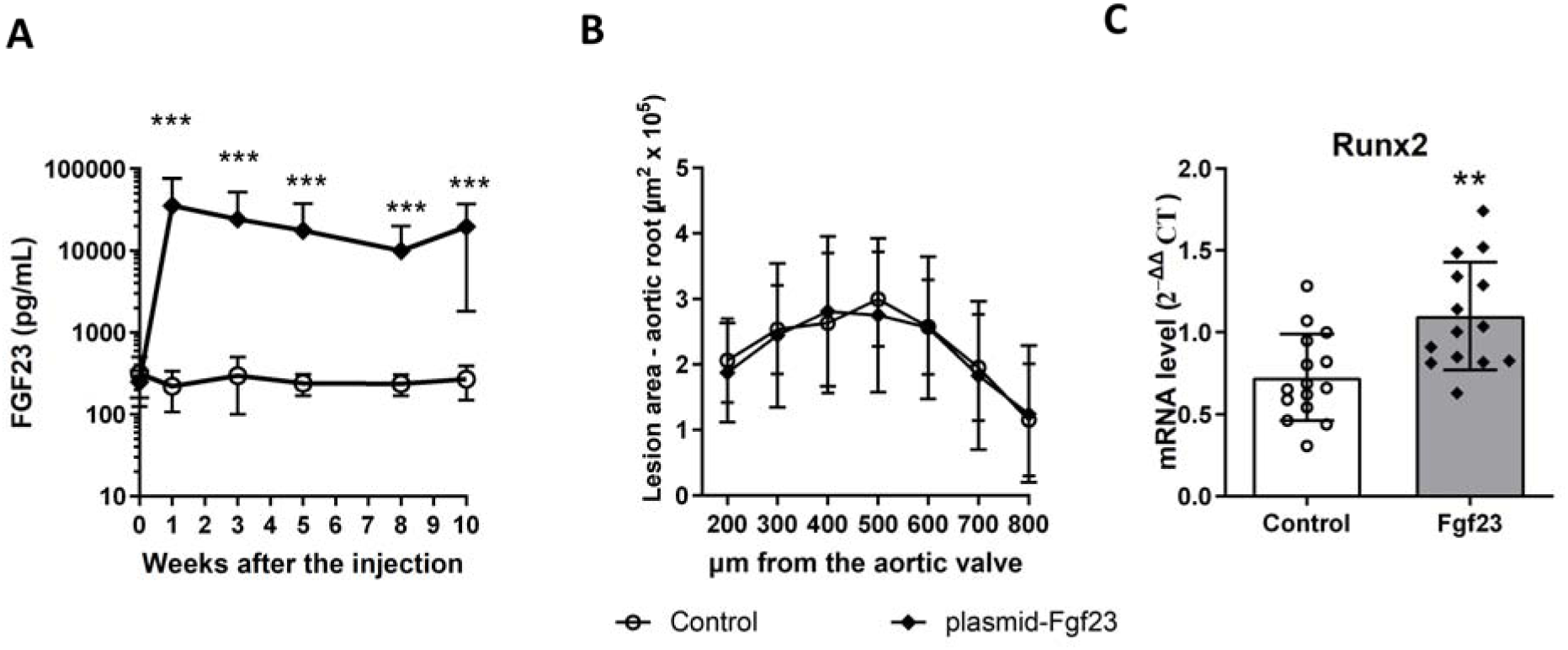
*ApoE-/-* female mice received a single hydrodynamics-based transfection of Fgf23-containing plasmid. Overexpression of FGF23 did not increase atherosclerotic burden in female mice. (A) Plasma FGF23 levels were analyzed at baseline (week 0), at weeks 1, 3, 5, and 8 after the injection, and at the endpoint (10 weeks after the injection) to monitor efficiency of Fgf23 expression. (B) Aortic roots were stained with Oil Red 0. The absolute lesion area was calculated at 7 levels above the aortic root. (C) mRNA levels of Runx2 analyzed by quantitative real-time RT-PCR. Values for individual mice are presented as 2-Mct value of a specific gene normalized to transcript level of HPRT, as a house-keeping gene. *\*\*P* < 0.01. n of mice= 15 and 13 for control and Fgf23 groups, respectively. Error bars indicate standard deviation.

**Supplemental Figure 3.**
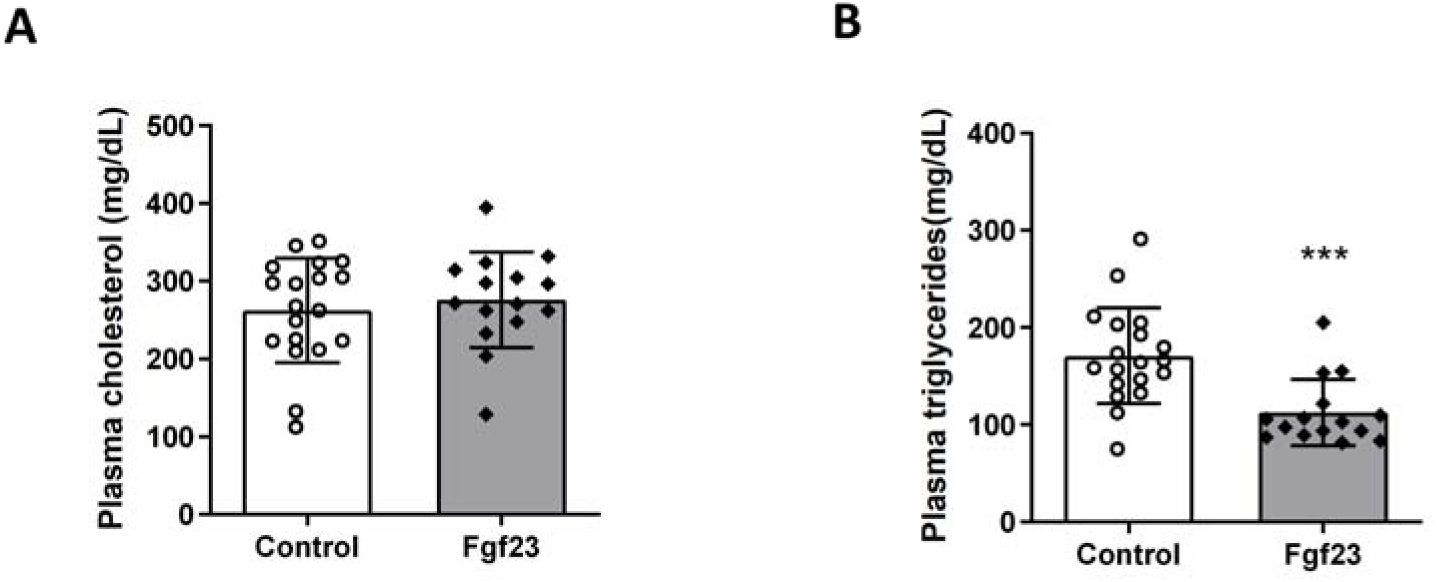
Lipid Analysis in *ApoE-/-* male mice. Levels of total cholesterol (A) and triglycerides (B) were measured in plasma. n = 19 and 15 for controls and for Fgf23 group, respectively. *** *P<0.001.* Error bars indicate standard deviation.

**Supplemental Figure 4:**
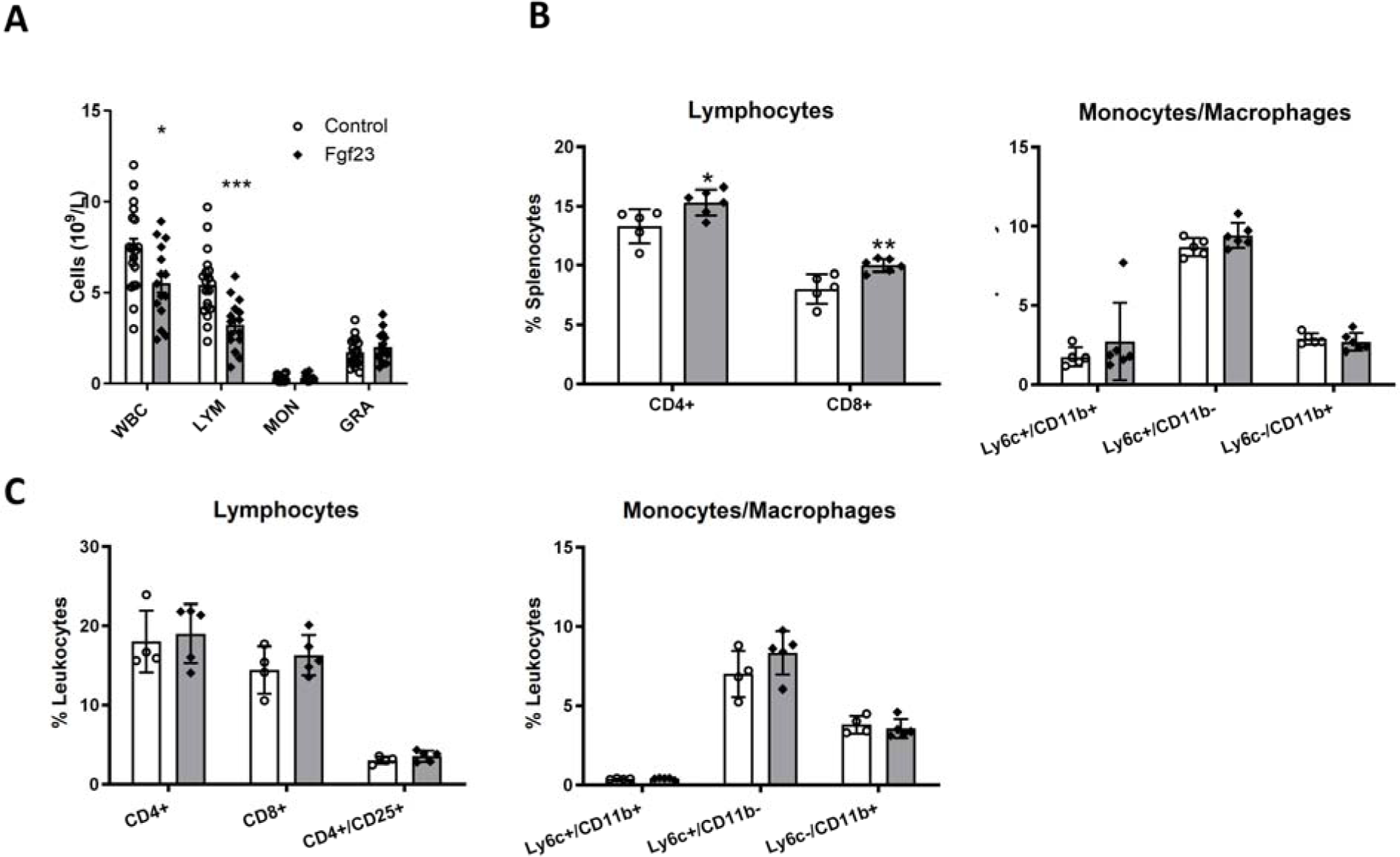
FGF23 overexpression leads to lymphocytopenia due to relocation of lymphocytes to spleen in *ApoE-/-* male mice. (A) Absolute numbers of total white blood cells (WBC) and separate populations of lymphocytes (LYM), monocytes (MON) and granulocytes (GRA) in the whole blood taken from mice at the endpoint of the experiment; n of mice= 19 and 15, for controls and Fgf23 groups, respectively; *** P<0.001 (B-C) Percentage of different subgroups of lymphocytes and monocyte/macrophages in spleens (B) and in paraaortic lymph nodes (C) taken from mice at the endpoint of the experiment; *\*P* < 0.05; *\*\*P* < 0.01, n = 5-6 per group. Error bars indicate standard deviation.

